# A spectral framework to map QTLs affecting joint differential networks of gene co-expression

**DOI:** 10.1101/2024.03.29.587398

**Authors:** Jiaxin Hu, Jesse N. Weber, Lauren E. Fuess, Natalie C. Steinel, Daniel I. Bolnick, Miaoyan Wang

**Affiliations:** Department of Statistics, University of Wisconsin-Madison; Department of Integrative Biology, University of Wisconsin-Madison; Department of Biology, Texas State University; Department of Biological Sciences, University of Massachusetts Lowell; Department of Ecology and Evolutionary Biology, University of Connecticut

**Keywords:** statistical genetics, gene co-expression network, expression quantitative trait loci mapping, network science, tensor methods

## Abstract

Studying the mechanisms underlying the genotype-phenotype association is crucial in genetics. Gene expression studies have deepened our understanding of the genotype → expression → phenotype mechanisms. However, traditional expression quantitative trait loci (eQTL) methods often overlook the critical role of gene co-expression networks in translating genotype into phenotype. This gap highlights the need for more powerful statistical methods to analyze genotype → network → phenotype mechanism. Here, we develop a network-based method, called snQTL, to map quantitative trait loci affecting gene co-expression networks. Our approach tests the association between genotypes and joint differential networks of gene co-expression via a tensor-based spectral statistics, thereby overcoming the ubiquitous multiple testing challenges in existing methods. We demonstrate the effectiveness of snQTL in the analysis of three-spined stickleback (*Gasterosteus aculeatus*) data. Compared to conventional methods, our method snQTL uncovers chromosomal regions affecting gene co-expression networks, including one strong candidate gene that would have been missed by traditional eQTL analyses. Our framework suggests the limitation of current approaches and offers a powerful network-based tool for functional loci discoveries.

**Significance statement:** This work addresses a key gap in understanding the mechanistic foundations for genotype-phenotype associations. While existing expression quantitative trait loci (eQTL) methods identify candidate loci affecting gene expression variants, they often neglect the crucial role of gene co-expression networks. Here, we develop a network-based QTL framework to map genetic loci affecting the gene co-expression network. Utilizing a tensor-based spectral approach, our snQTL method estimates the differential co-expression patterns and effectively identifies the associated genetic loci. Application of snQTL to three-spined sticklebacks revealed candidate loci missed by standard methods. This work suggests the limitations of current approaches and highlights the potential of network-based functional loci discovery.

## 1 Introduction

The identification of genetic variants underlying complex phenotypic traits has been a pivotal area in genetics research for decades. Genome-wide association studies (GWASs) have identified important genetic variants by detecting statistical association between phenotypes and genotypes in outbred populations [31]. Likewise, quantitative trait locus (QTL) mapping in experimentally crossbred organisms allows researchers to shuffle genetic backgrounds meiotically and test for associations between measurable phenotypes and chromosomal regions. However, both GWAS and QTL mapping are limited by the challenge of elucidating the mechanisms behind these genotype-phenotype associations, and the lack of sufficient functional information for many loci [45, 40]. Gene expression studies can bridge this gap between genotype and phenotype. To this end, expression quantitative trait locus (eQTL) analysis was developed to identify associations between genetic variants and gene expression levels [18]. The eQTL studies have deepened our understanding of genotype → expression → phenotype mechanisms [16, 23, 45, 43]. Existing eQTL methods have identified numerous genetic loci, categorized as cis- or trans-eQTL, that influence gene expression. The cis-eQTLs are located near the expressed gene on the same chromosome, and they typically directly affect the binding of transcription factors or chromatin proteins to DNA [6, 10]. Conversely, trans-eQTLs reside on different chromosomes and often influence the expression or structure of transcription factors, ultimately impacting their ability to regulate the expression of distant genes [29, 37, 32].

A key limitation of current eQTL studies is their focus on individual genes, but not on the network structure of gene co-expression. Gene co-expression networks are often represented by correlation matrices at the whole-transcriptome scale [27, 25]. Correlation among gene expressions may arise, for example, when multiple genes are co-regulated by the same transcription factor or participate in sequential regulatory cascades. Correlated expression can also arise from genetic linkage between separate regulatory cascades and from shared environmental effects, although in this case correlated expression do not necessarily imply direct functional interactions.

There is accumulating evidence that gene co-expression networks can differ between species [20, 5] or populations [20], even in controlled environments. These differences suggest the gene co-expression network is evolvable, and hence most likely has a genetic basis. The genetic-related co-expression might occur, for instance, if one allele of transcription factor A controls expression of genes B and C (creating correlations between A, B, and C); but the alternate allele controls only gene B but not C (eliminating the AC and BC correlation). Mutations that alter gene linkage patterns (e.g., inversions or translocations) could also alter gene co-expression networks. This concept is similar to mapping epistatic eQTLs [11], except that those studies (excluding work on highly prolific laboratory models) only rarely have the power to identify more than a few interacting genes. If genetic variants broadly alter co-expression network structure, eQTL or GWAS methods could in principle map these genetic loci. Analyzing these associations between genetic loci and co-expression network can reveal the network-level impact of quantitative trait loci, leading to new insights into the genetic basis of complex traits. Developing efficient methods for network-based eQTL is a topic of great interest.

Recent studies have extended the concept of eQTL to co-QTLs [30, 1, 40, 14]. These methods aim to identify genetic loci that explain coordinated changes in expression between pairs of genes. However, current co-QTL methods have several limitations. One of the challenges is the massive number of statistical tests needed, which increases exponentially with the number of gene pairs analyzed. Some methods restrict co-QTL searches to previously identified eQTLs [30, 14], while others prioritize gene pairs based on prior knowledge [40]. These approaches reduce testing burdens but may miss important co-QTLs. Furthermore, current co-QTL methods are limited by assuming linear models with additive effects [30, 1, 40, 14]. The additive assumption neglects dominance, recessiveness, or even transgressive inheritance, hindering the ability to capture the full genetic influence on co-expression networks. More powerful network-based eQTL methods are needed to address these issues.

In this paper, we propose a novel method called spectral network QTL (snQTL) to address these challenges. Our snQTL identifies the association between genotype and the entire co-expression network structure. The snQTL method identifies network-QTLs (nQTLs) which explain a fraction of the genetic variance of the entire gene co-expression network. The snQTL represent genetic variants that alter the global pattern of a network, while traditional co-QTLs represent genetic variants that alter the expression for only a particular pair of genes. Statistically, the key idea of the snQTL method is to use tensor spectral statistics to represent the joint difference in gene co-expression networks at each of many different loci. This approach reduces the number of tests to the number of genetic markers throughout the genome of a recombinant hybrid population (used for mapping), and we allow for the simultaneous consideration of all active genes in the network. We also propose a permutation-based approach to obtain valid testing results that are robust to the data distribution. In addition to identifying nQTLs, our snQTL framework also outputs the joint differential networks, which represents the specific network patterns that are altered by genetic variants at the detected nQTLs. Our approach has the potential to be extended to mapping genetic effects on the architecture of microbiome co-occurrence networks and proteomic networks. We demonstrate the effectiveness of our method in the immune tissue gene expression data from a large genetic cross of three-spined stickleback fish (*Gasterosteus aculeatus*).

## 2 Results

### 2.1 Spectral network QTL framework

Figure 1 illustrates the main framework of our snQTL method. We take as input (i) expression read counts of *p* genes and (ii) genotypes of *m* genetic markers, from the same set of *n* individuals. The snQTL method then outputs two key results at each marker: (i) a p-value indicating the association significance between the co-expression network and the marker, and (ii) a joint differential network with nodes representing genes and edges representing associated effects. The individuals in the study are drawn from an F2 hybrid generation derived from crosses between genetically divergent populations, with parent-of-origin diagnostic genetic markers spread across all chromosomes.

**Figure 1:**
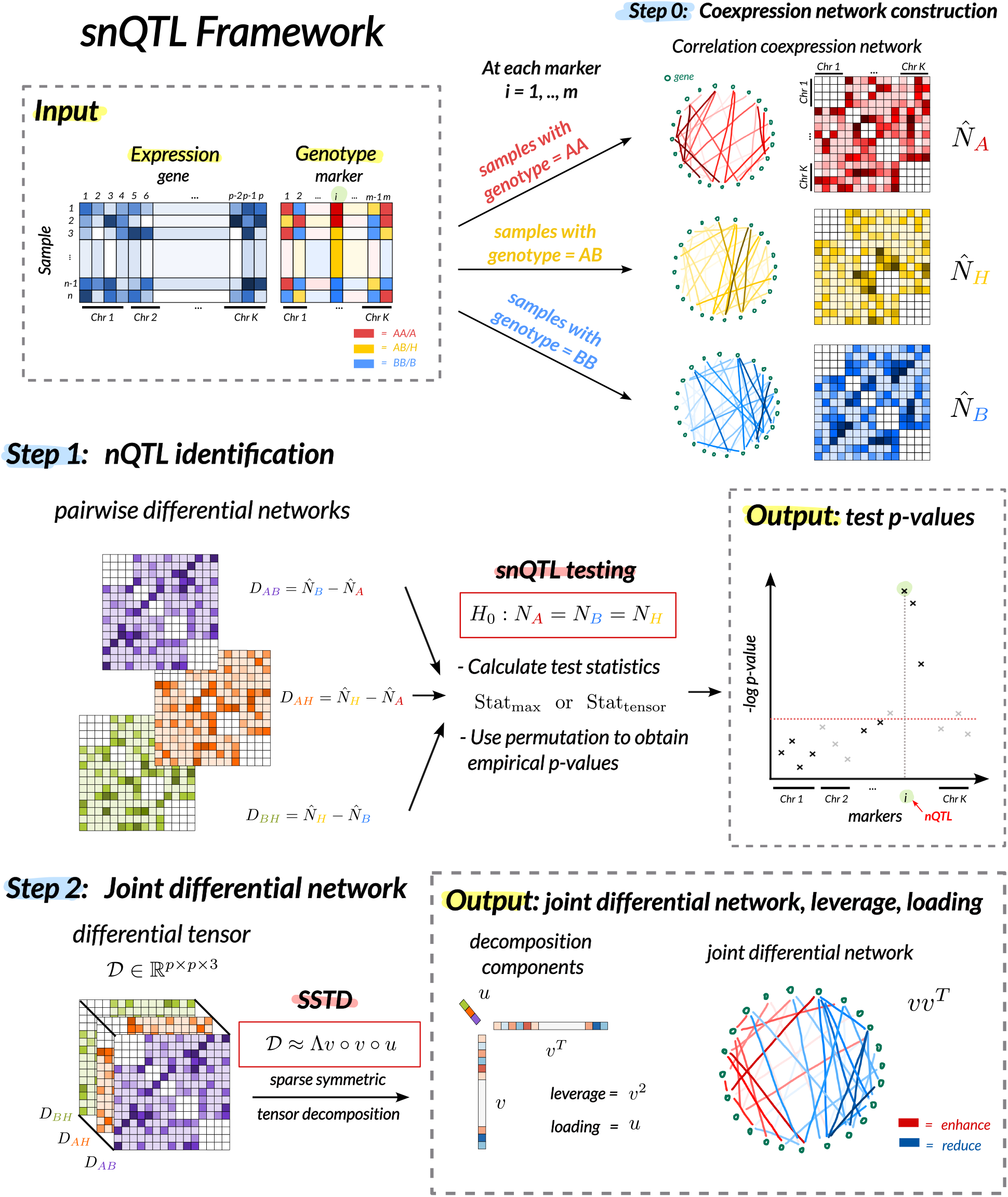
The main idea of our snQTL framework. Our snQTL framework takes as input (i) gene expression read counts and (ii) genotypes of genetic markers from the same set of samples. The snQTL consists of three steps: (0) co-expression network construction, (1) nQTL identification via hypothesis testing using multilinear spectral statistics, and (2) joint differential network estimation at associated loci via sparse symmetric tensor decomposition. At each marker, the output includes (i) a p-value indicating the association significance between the co-expression network and the marker, and (ii) a joint differential network with nodes representing genes and edges representing associated effects.

The snQTL consists of three steps. First, we construct gene co-expression networks; see Step 0 in Figure 1. At each of the *m* markers, we split the gene expression data by genotype (AA, AB, BB) and calculate Pearson correlation matrices within each group. Let *N*_*A*_, *N*_*B*_, *N*_*H*_ denote the (unknown) population correlation matrices, where *A* and *B* denote homozygous genotypes and *H* denotes heterozygous genotype. Because linkage disequilibrium (LD) in F2 hybrid crosses often causes strong correlations between genes on the same chromosome, the co-expression networks will be enriched for non-functional within-chromosome edges. To focus on trans-snQTL effects, we exclude within-chromosome correlations by setting the (*j, k*)-th entries in *N*_*A*_, *N*_*B*_, *N*_*H*_ to zero, if genes *j* and *k* are located on the same chromosome.

Next, we perform statistical tests to identify genetic markers affecting co-expression networks; see Step 1 in Figure 1. At each marker *i*, we test the null hypothesis:

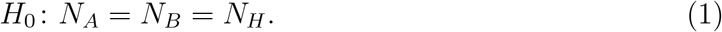

In the next section, we will provide several test statistics based on the multilinear spectral components of the correlation matrices. Let 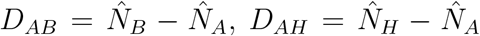, and 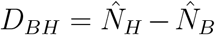 denote the three pairwise differential networks, where 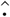 denotes the sample correlation matrix. The multilinear spectral components of differential networks allow us to test for classical genetic dominance effects as well as a broad range of genetic effects onto the entire co-expression networks. We use permutation to obtain the *p*-value for the hypothesis test in (1). The output is summarized as a Manhattan plot of association *p*-values across the genome.

Last, we estimate the joint differential network at the associated marker; see Step 2 in Figure 1. We use sparse tensor decomposition to obtain the leading eigenvectors in the pairwise differential correlations. These eigenvectors summarize the differential signal into a single network. The resulting joint differential network has nodes representing genes and edges representing co-expression changes associated with the genetic marker.

#### snQTL testing and joint differential network estimation via sparse tensor decomposition

We briefly introduce the sparse symmetric tensor decomposition (SSTD) in our contexts. Let 𝒟 ∈ ℝ^*p*×*p*×*q*^ be an order-3 tensor with each of the *q* slides being a symmetric *p*-by-*p* matrix. We say 𝒟 is sparse and of rank 1 if 𝒟 satisfies the SSTD model:

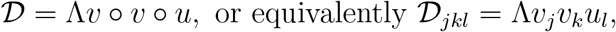

for all (*j, k, l*) ∈ {1, …, *p*} × {1, …, *p*} × {1, …, *q*}, where *°* denotes the vector outer product, *v* and *u* are norm-1 vectors in ℝ^*p*^ and ℝ^*q*^, respectively, and *v* is further sparse with ∥*v*∥_0_ ≤ *R* for some constant *R* ≤ *p*, and Λ ∈ ℝ_+_. Here ∥ · ∥_0_ is the L0 norm that counts the number of non-zero entries in the vector. The constraint on ∥*v*∥_0_ controls the sparsity on the first two modes. We call Λ, *v*, and *u*, the sparse leading tensor eigenvalue (sLTE), the sparse tensor eigenvector, and the loading vector, respectively.

In our snQTL framework, we define an order-3 differential tensor 𝒟 ∈ ℝ^*p*×*p*×3^ by stacking the three pairwise differential networks *D*_*AB*_, *D*_*AH*_, *D*_*BH*_ together. To summarize the signal in 𝒟, we compute the SSTD approximation to the tensor 𝒟. Specifically, we solve for the spectral components (Λ, *v, u*) that minimize the least square approximation error

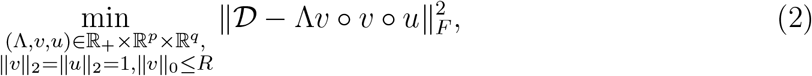

where ∥ · ∥_*F*_ denotes the Frobenius norm defined as the squared sum of tensor entries, and ∥ · ∥_2_ denotes the vector L2 norm. We denote the sLTE solution as Λ(𝒟), with 𝒟 being the input differential tensor. A larger sLTE suggests a stronger signal against the null in (1).

Our test statistics, named Stat_test_, is defined using the sLTE:

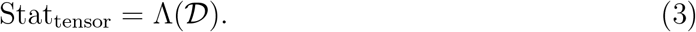

Our snQTL also features the estimation of a joint differential network. The sparse tensor eigenvector, *v* = *v*(𝒟), and loading vector, *u* = *u*(𝒟), together capture a lower-dimensional representation of 𝒟. We call the leading matrix approximation, *v*(𝒟)*v*(𝒟)^*T*^, the “joint differential network”. This network captures the overall co-expression network changes in response to the genetic variation at the marker of interest. We call the element-wise squared eigenvector, denoted as *v*^2^, the “gene leverage”. The leverage represents the overall connectivity of node (i.e. gene) in the joint differential network. Genes with higher leverage are highly connected in the joint differential network and thus contribute more to the overall differential signal (as captured by the sLTE in (3)). The loading vector, *u*(𝒟), reveals the weights of pairwise comparisons towards the joint differential network. Higher values in magnitude in *u*(𝒟) represent a higher contribution of the corresponding pairwise comparison towards the joint comparison.

Our snQTL is inspired from earlier work on SSTD [28]. However, we introduce key modifications that tailor SSTD to our specific needs in snQTL analysis. We explicitly considers the symmetry and sparsity in the first two modes of the tensor, making SSTD a better fit for our framework (details in *Materials and Methods*). Furthermore, unlike earlier work that focuses on the decomposition only [28], our primary goal is hypothesis testing within the context of snQTL analysis. We have developed specific tools for this purpose.

#### snQTL testing via sparse matrix decomposition

We also propose an optional statistic for (1), based on extension of sparse leading matrix eigenvalue (sLME) [44]. The sLME of a matrix *D* is defined as

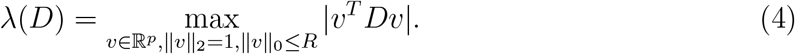

The sLME represents the maximum eigenvalue of matrix *D* subject to the sparse eigenvectors. Our second test statistics, named “max”, is defined as the maximal sLME from all three pairwise comparisons:

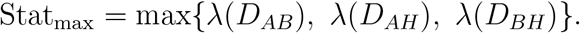

Under the null hypothesis in (1), all pairwise differences (*D*_*AB*_, *D*_*AH*_, *D*_*BH*_) are zero matrices, resulting in a zero max statistic. Conversely, a larger max statistic indicates higher differences in at least one pairwise comparison, making it well-suited for joint comparison of multiple networks.

Our max statistic generalizes the earlier work from pairwise comparison [44] to joint comparison of multiple matrices Other methods include *L*_2_-type statistics [13] that consider all entries in the comparison, and *L*_*∞*_-type statistics [2] that focus on the largest deviation. However, the *L*_2_-type statistics assume all genes contribute equally, while the *L*_*∞*_-type statistics capture only the single most extreme gene pair. In contrast, the spectral statistic, sLME, is well-suited for scenarios where the signal is weak and sparse, meaning that a small subset of genes exhibit moderate effects. This aligns with the biological expectation that genes might have significant but subtle co-expression changes. Additionally, the sparsity in sMLE promotes result interpretability and faster computation.

#### Algorithm implementation

We design an iterative algorithm that alternatively updates the decomposition components to approximately solve (2). We adopt the penalized matrix decomposition [44, 38] to approximately solve for sLME in (4). In practice, we also consider variants of tensor and max statistics, such as the sum of sLMEs and the squared sLTE (*Supporting Information Text*). More variants can be designed based on problem contexts. For all test statistics, we use permutation to approximate the null distributions and obtain the empirical p-values. The number of permutations and the sparsity hyper-parameter *R* can be adjusted as needed. See *Materials and Methods* for more details.

#### Analysis of simulated data

We first evaluated the efficiency of our snQTL framework on synthetic data for 200 genes across 20 chromosomes. We started with genetically divergent homozygous parents, and simulated the genotypes for an F1 cross and for an F2 intercross generation with random chromosomal crossing overs. For each F1 gamete, we simulated one recombination event per chromosome per gamete, randomly placed along the chromosome with a uniform distribution. The F2 hybrids’ gene expression counts were generated from Poisson distributions with parameters varying by genotypes. We randomly selected one gene as the nQTL and altered the expressions of target genes based on the additive network effect associated with the genotype at the selected nQTL. Note that we took all 200 genes as the candidate loci for nQTL in the simulations. Sets of other genetic markers such as single nucleotide polymorphisms may serve as the suitable set of candidate loci for nQTL identification under real scenarios. We tested the framework with varying hybrid population sizes from 50 to 500 to assess performance cover various scenarios.

Figure 2 confirms the similarity between the synthetic and real F2 hybrid three-spined stickleback data [35]. The similar block diagonal patterns in the genetic correlation heatmaps (Figure 2A) suggest the LD among real and simulated markers. The overlapped histograms of expression counts (Figure 2B) validate our simulation procedures, indicating parameter values effectively mimicked real datasets.

**Figure 2:**
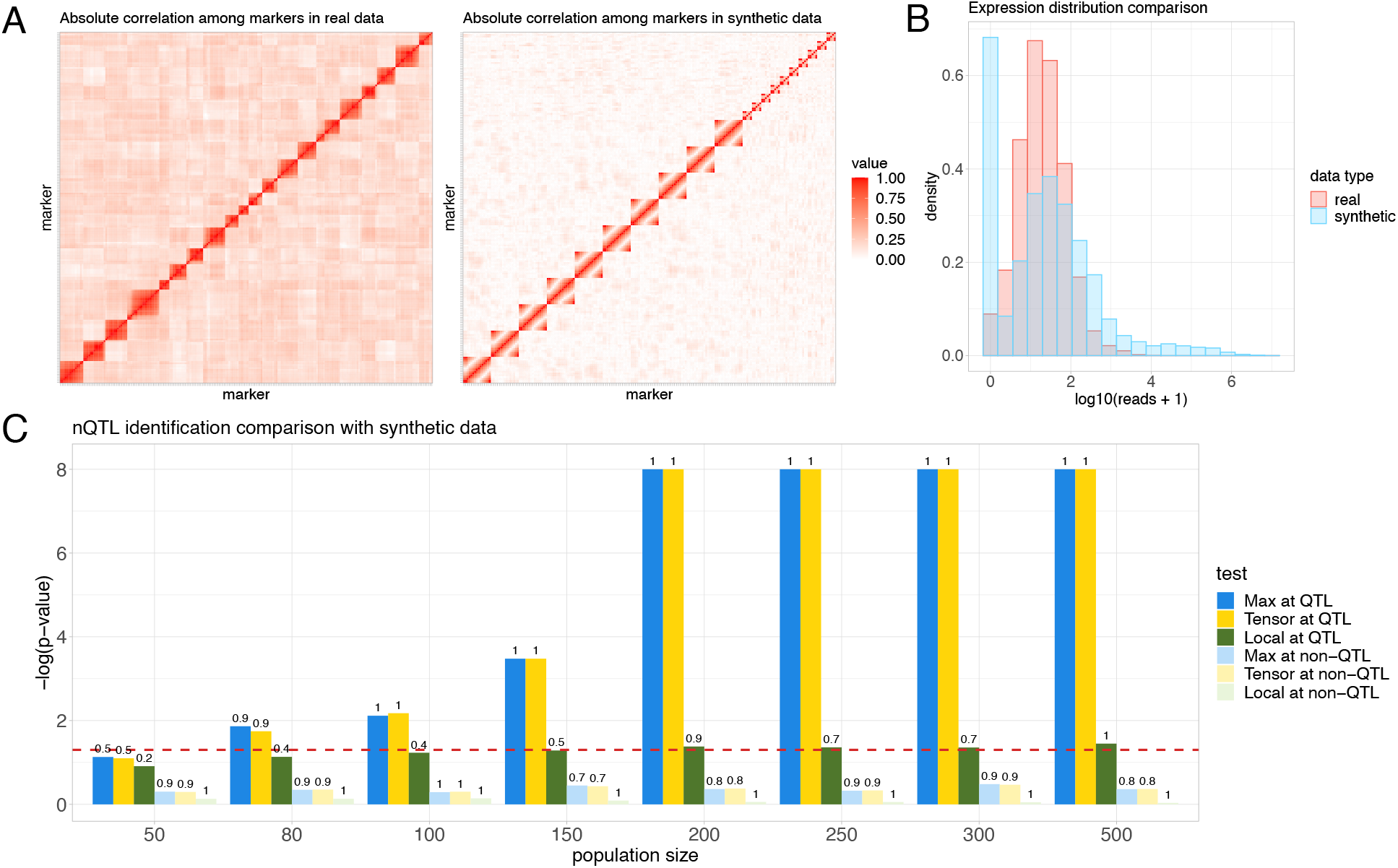
Analysis of simulated data. Synthetic datasets in three panels have the same parameter setup. (A) Absolute genetic correlation heatmaps among the markers in real F2 hybrid three-spined stickleback data [35] and synthetic data. Markers are ordered following their positions on the genome. Genetic correlations are measured by absolute sample Pearson correlation coefficients between the genotypes of two markers. (B) Density histograms for expression counts in real stickleback and synthetic data. (C) Barplots comparing the nQTL identification performances for snQTL framework and local method (F-test for regression of pairwise co-expression onto genotype) on synthetic data with varying population size from 50 to 500. Red dashed line corresponds to the critical threshold of p-value 0.05. True positive (or negative) rates for the tests at nQTL (or non-nQTL) are shown above the bars. All reported numbers are averaged across 15 replications for each population size.

We compared three methods on the synthetic data: the snQTL framework with max statistic, with tensor statistic, and a local approach based on F-tests for linear regressions of pariwise co-expression against genotypes. This local approach is similar to previous co-QTL analyses [30, 40]. We assessed both statistical power and type I error by applying all tests at the nQTL and non-nQTLs. Average test p-values and true positive (TP)/negative (TN) rates were recorded across 15 replicates for each population size.

Figure 2C demonstrates the superior statistical power of our snQTL framework, especially with larger populations. The out-performance suggests that the snQTL framework effectively addresses the multiple testing burden and tends to lead to more discoveries than the local approach. Additionally, the high TN rates at non-nQTLs support the high accuracy of the snQTL framework for nQTL identification.

### 2.2 Performing snQTL to map stickleback loci affecting co-expression networks

We conducted snQTL analysis on the three-spined stickleback (*Gasterosteus aculeatus*) data [35] to reveal the genetic landscape for co-expression networks in sticklebacks. These datasets are from a QTL mapping study in which wild fish were obtained from two lakes on Vancouver Island (Roberts Lake and Gosling Lake; RR and GG), and eggs/sperm mixed in petri dishes to generate F1 hybrids (RG). These hybrids were reared to maturity in an aquarium lab at the University of Texas and intercrossed to generate F2 intercross hybrids (RG*RG) and reciprocal backcrosses (RG*GG, GG*RG, RG*RR, RR*RG). Although hybrid crosses constituted a mixture of maternal backgrounds, maternal effects were excluded in our analyses. All F2 generation fish were reared to maturity in the laboratory and experimentally exposed to a cestode parasite, then euthanized 42 days post-exposure. Transcriptomic dataset was collected from head kidneys (pronephros, a major immune organ in fish) using Tag-Seq [15]. The cross design, sequencing methods, and bioinformatics pipelines are described in depth in earlier work [35, 9].

The raw dataset consists of gene transcript counts and genotypes for 234 markers, for 351 samples from F2 generations and backcrosses. We preprocessed the data with the following procedure. First, to eliminate non-functional variations, we normalized the read count matrix and regressed expressions against the sex and ancestry covariates, retaining the residuals (*Supporting Information Text*). Second, we focused the analysis on the top 10,000 genes with the highest adjusted mean expressions, as more information may be involved with actively highly expressed genes. The cutoff of 10,000 was chosen to ensure computational efficiency.

In addition, we considered the infection status of the sample fish as cestode infection is likely an environmental confounder. We added the worm presence as a predictor in the pre-processing regression step. Our snQTL analysis exhibited the same conclusions (*Supporting Information Text*) before and after the additional procedure, suggesting the robustness of our discoveries to the infection status. For conciseness, we presented only the analysis without infection covariates in this paper. We leave further analyses involving more covariates and genes for future investigations.

#### Identification of stickleback nQTLs

We performed snQTL analysis on stickleback data using both tensor and max statistics. Both approaches lead to similar testing results (*Supporting Information Text*), demonstrating the robustness of our nQTL identification. e present the findings using the tensor statistic here, as the tensor approach also facilitates joint differential network estimation. The Manhattan plot in Figure 3A shows 21 stickleback nQTLs concentrated at Chr 3, Chr 8, and Chr 18. This clustering pattern of nQTLs aligns with the LD structure among markers (Figure 2A). The three chromosomes of interest all exhibit extensive and stronger signals of snQTL associations compared to other chromosomes. To further narrow down potential functional regions, we examined within each snQTL region for coding genes with strong genomic signatures of past natural selection. Specifically, we used published population genomic data: allele frequency estimates obtained from PoolSeq of *∼* 100 fish from each of three populations (Roberts Lake, Gosling Lake, and a marine outgroup). We calculated population branch statistics (PBS) measuring accelerated evolution in each lake (Roberts or Gosling), relative to an ancestral marine population (Sayward), as described in earlier work [35]. Large PBS in either lake population indicates a gene that was likely a target of natural selection within the lake in question, since its colonization *∼* 12,000 years ago.

**Figure 3:**
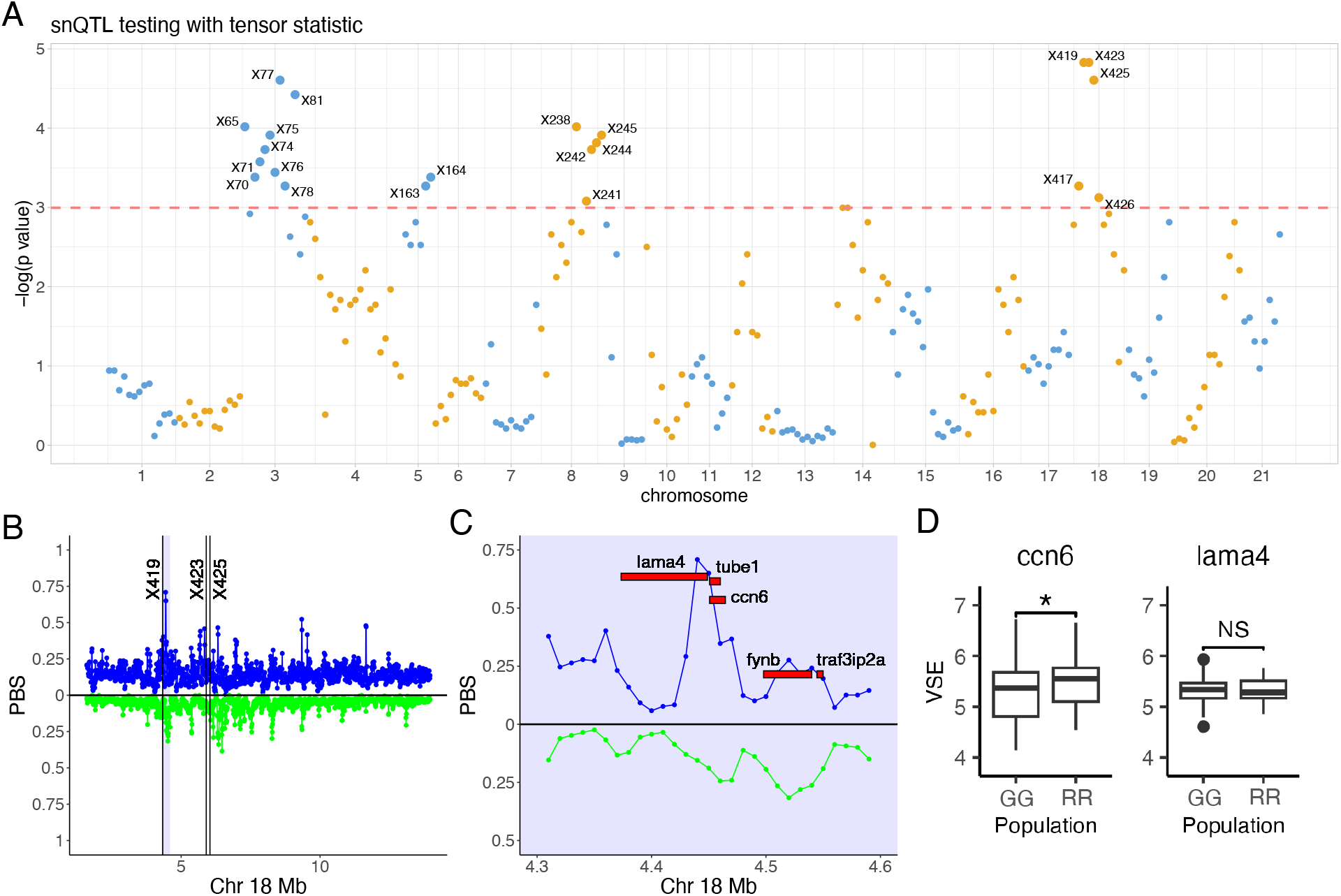
Identification of stickleback nQTLs via snQTL framework. testing with tensor statistics marks 21 stickleback nQTLs (above pink dashed line, with p-values smaller than 0.05), mainly clustered in Chr 3, Chr 8, and Chr 18. (B) Strong genomic targets of selection with high population branch statistic (PBS) distribute around the outstanding nQTLs (markers X419, X423, and X425) in Chr 18. Values above the medial line represent higher PBS in Gosling Lake (blue); values below the line represent higher PBS in Roberts Lake (green). (C) Zoomed-in shadowed area in (B). Development regulation genes, lama4 an d ccn6, locate tightly around marker X419 with high selection speed. (D) Variance stabilized expressions (VSE) for ccn6 and lama4 in Gosling (GG) and Roberts (RR) lakes.

Several protein-coding genes lie in regions adjacent to PBS outliers within nQTLs (*Supporting Information Text*). We focused our analysis on genes near the largest nQTL on Chr 18 (Figure 3). None of these genes harbored coding variants but two were represented in our expression data: cellular communication network factor 6 (*ccn6*) and laminin subunity alpha 4 (*Lama4*). Although the *Lama4* expression differs little between parental populations, the *ccn6* expression was significantly lower in Gosling fish (*t* = 2.115, *df* = 97.886, *p* = 0.037, Figure 3D). The gene *Ccn6*, also known as *wisp3*, has 4 distinct protein domains that perform distinct functions [22], several of which have notable connections to the stickleback system. Secreted *ccn6* can bind to and limit insulin growth factor-1 (*igf-1*) signaling, thereby suppressing cell growth and metabolic potential [24], as well as mediating fibrotic responses [39, 26]. The gene *Ccn6* also acts as a transcription factor that activates genes necessary for formation of the mitochondrial electron transport system [21] and indirectly regulates reactive oxygen species (ROS) levels [17]. Gosling fish produce significantly less ROS, display less cestode-induced fibrosis, and grow faster than Roberts fish. It is worthy noting that in humans, the *ccn6* expression is largely restricted to kidney, skin and testes, consistent with an organ-specific regulatory role [35, 7].

#### Joint differential network at nQTL locus X419

We further estimated joint differential networks for the significant nQTLs identified in our snQTL analysis. We found that most nQTLs are associated with similar sets of genes with high leverages, resulting in joint differential networks with comparable patterns (*Supporting Information Text*). The result suggests that our snQTL approach captured the robust and global co-expression patterns associated with genetic variation.

Here, we present the joint differential network at the most significant nQTL, X419 on Chr 18. We ranked genes based on their leverage scores from our method. We found that the top 10 genes achieved a cumulative leverage of 0.54, and the top 100 genes achieved a cumulative leverage of 0.9. We called the top 10 genes with highest leverages the “primary genes”, and the remaining top genes the “secondary genes”. These top 100 genes distribute widely on the genome, from the scaffold region and mitochondrial genome (MT) to all chromosomes (Figure 4A). This wide distribution of top genes implies the capacity of nQTLs to impact co-expressions throughout the whole genome. Such cross-chromosome influences are likely to represent functional genotype-network associations. In addition, the loading values for the genotype comparisons GG-RG and RG-RR are 0.498 and 0.31, respectively. The result suggests that the co-expressions between primary and secondary genes, except those with *bbx* and *otog*, are reduced in Gosling Lake fish and enhanced in Roberts Lake fish (Figure 4B). Moreover, the loading and co-expression networks for three genotypes (Figure 4C-E) show that differential networks for GG-RG and for GG-RG are comparable, indicating the nearly additive genetic effects to the co-expression networks.

**Figure 4:**
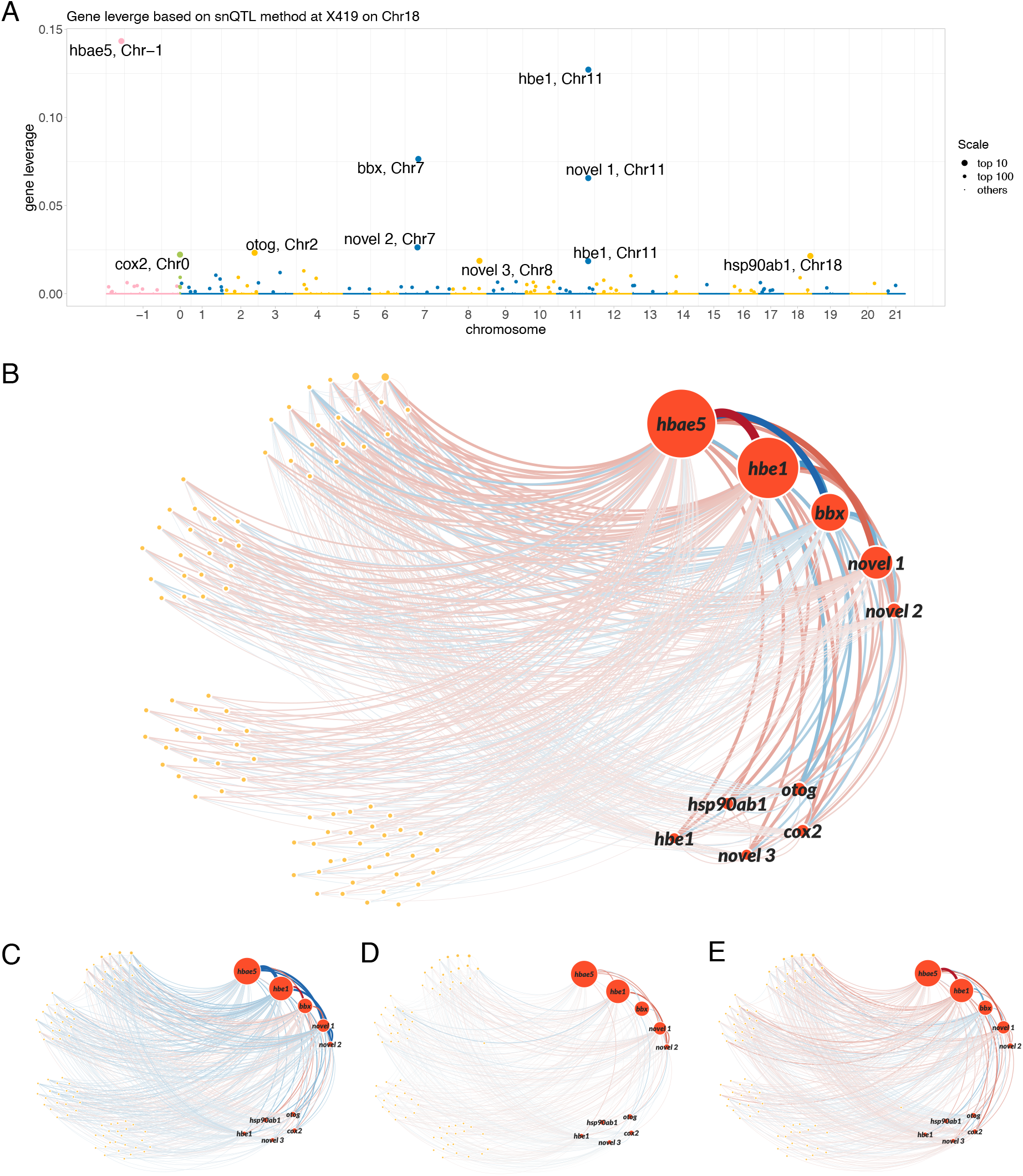
Joint differential network analysis at nQTL X419 on Chr 18. (A) Leverage scores for 10000 genes. Primary genes with top 10 leverage are highlighted with transcription IDs. Mitochondrial genome (MT) and scaffold region are coded as Chr 0 and Chr −1, respectively. (B-E) Networks for primary (red annotated nodes) and secondary (orange nodes) genes with top 100 leverages. The edge width indicates the connection strength between two genes; the diameter of node indicates the leverage of the gene; the color indicates enhancement (red) or reduction (blue) of the connection compared with average level. (B) Joint differential network at X419 with top 10% strongly connected edges. A wider edge implies a stronger genetic variation in the co-expression of the gene pair. Most genetic co-expression variations occur between the primary and secondary genes. (C-E) co-expression networks corresponding to the genotypes GG, RG, and RR at X419, respectively. The linear changes in the colors of edges imply the nearly additive genetic effect to the co-expression networks. novel 1: ENSGACT00000018413; novel 2: ENSGACT00000026589; novel 3: ENSGACT00000017116.

We found that most genetic co-expression variations occur between the primary and secondary genes (Figure 4B). We note that many of the primary genes (*hbae5*, two *hbe1* paralogs, and the novel gene *ENSGACT00000018413*, which is orthologous to *hba2* in other species of fish) are hemoglobin subunits expressed in red blood cells and directly participate in oxygen transport activities, while the others are involved in closely related biological processes, such as blood vessel development (*hsp90ab1*) and carbohydrate metabolism (*otog*) (Table 1). These functions are consistent with decreased expression of *ccn6* being connected to elevated rates of *igf-1* signaling and cell replication in the head kidney, which is the hematopoietic organ in fish. Similarly, overexpression of heat shock proteins (i.e., *hsp90ab1*) can be stimulated either via pharmacological suppression of *igf-1* [33] or dysregulation of the electron transport chain in mitochondria, which is another major function of *ccn6*. Although the precise role of *mmp16b* has not been well characterized, *igf-1* is connected to the expression of other *mmp*s. Our analysis demonstrates the power of snQTL framework with functional annotation for unraveling the genetic basis of co-expression networks.

**Table 1:** List of primary genes with top 10 leverage scores in joint differential network at X419 on Chr 18.

| Transcript ID | Leverage | Gene | Chr | Gene and Protein GO annotations |
| --- | --- | --- | --- | --- |
| ENSGACT00000019169 | 0.1433 | hbae5 | Scaffold 112 | heme, iron ion, oxygen binding; oxygen carrier activity |
| ENSGACT00000018425 | 0.1271 | hbe1 | 11 | heme, iron ion, oxygen binding; oxygen carrier activity |
| ENSGACT00000026622 | 0.0765 | bbx | 7 | DNA binding |
| ENSGACT00000018413 | 0.0656 | - | 11 | heme, iron ion, oxygen binding; oxygen carrier activity |
| ENSGACT00000026589 | 0.0263 | - | 7 | uncharacterized |
| ENSGACT00000022959 | 0.0232 | otog | 2 | carbohydrate metabolic process |
| ENSGACT00000027730 | 0.0222 | cox2 | MT | copper ion binding; cytochrome-c oxidase activity in mitochondrion & respirasome |
| ENSGACT00000017921 | 0.0214 | hsp90ab1 | 18 | blood vessel development; leukocyte migration; response to estrogen |
| ENSGACT00000017116 | 0.0187 | - | 8 | serine-type endopeptidase activity; proteolysis |
| ENSGACT00000018389 | 0.0185 | hbe1 | 11 | heme, iron ion, oxygen binding; oxygen carrier activity |

## 3 Discussion

Gene co-expression networks play a pivotal role in translating genotype into phenotype. This suggests that phenotypic evolution may often be a consequence of evolution not just of single genes’ protein structure or expression level, but also by changes of co-expression patterns among genes [19, 20, 5]. For gene co-expression networks to evolve, there must be genetic variations within species that impact the network structure, which selection (or drift) might act on. Therefore, there is a need for methods capable of identifying loci (or chromosomal regions) that are associated with changes in co-expression networks. While methods exist for analysing pairwise gene co-expression [30, 1, 40, 14, 36, 12], a key challenge lies in methods that can analyze gene co-expression across entire networks.

### 3.1 Methodological significance

Our snQTL framework offers a methodological advance in network-based association study. Unlike traditional co-QTL methods that test millions of gene pairs independently, snQTL treats the entire co-expression network as a single entity. This dramatically reduces the multiple testing burden. Furthermore, snQTL leverages a tensor spectral statistic that captures the overall signal across the entire network. This approach avoids the need for pre-selecting candidate gene pairs, which can introduce bias. Additionally, unlike regression-based methods that assume an additive genetic effect, snQTL allows for a broad range of genetic effects. The flexibility enables the detection of nQTLs as long as a significant difference exsits in co-expression network between genotypes.

The power of snQTL extends beyond co-expression networks. The framework can be generalized to analyze various networks, including microbial networks, proteomic networks, and others. With minor adjustments, snQTL can also handle directed networks like transcription factor binding networks and metabolic network. The core idea of snQTL can be applied for general mapping tasks beyond genetics. For example, the method can handle comparisons of more than three networks, alloqing investigation of associations with various discrete factors, such as treatment, location, or environmental conditions.

Several future improvement can be made to snQTL. Currently, snQTL removes all within-chromosome co-expression to address LD. Future improvements could incorporate recombination maps to identify unlinked markers on the same chromosome and linked markers on different chromosomes, providing a more biologically relevant approach. The other potential extension is on the use of SSTD. The current rank-1 SSTD approximation in snQTL captures the strongest signal in the network difference. Extending this to a higher-rank model could reveal more delicate signals, potentially leading to additional discoveries.

### 3.2 Biological significance

One of the “grand challenges” of biology is to understand the details of how genotypes produce phenotypes, and thereby develop tools to predict phenotypes. Genotype-phenotype prediction remains a challenge because most phenotypes are the emergent result of complex interactions between numerous genes. Network analyses offer a promising toolkit for representing these complex interactions. Such tools have been applied to gene-gene co-expression data [41, 42, 8], single-cell RNAseq data [34], gene-gene epistasis effects [4], proteomic data [3], and beyond, with the goal of describing the logic of genetic regulatory “circuits”. The hope is that this network-based approach can reveal rules of life not visible for single genes and their mRNA and protein products, or simple pairwise gene interactions.

Our snQTL analysis of three-spined stickleback gene expression illustrates this potential benefit. We identified three chromosomes with significant nQTLs. Using population genomic data, we were able to identify a candidate gene under especially strong selection within the nQTLs on Chr 18. The gene *ccn6* is a highly pleiotropic gene known to affect growth, metabolism, fibrosis, ROS production, and hence with great potential for network-wide effects in the immune organ sampled for transcriptomics. It appears likely that *ccn6* -mediated changes in electron transport chain function is affecting ROS production differences previously documented between the hybridized populations, with additional consequences for a protective fibrosis phenotype. This gene was not flagged in prior differential expression analyses of the same dataset. Although *ccn6* is expressed at significantly lower levels in Gosling than Roberts Lake fish, the differential expression was not exceptionally large. In contrast, the snQTL (aided by selection scans) makes this gene an important candidate for multivariate phenotypic effects. This result highlights a major limitation in how we currently search for expression-related evolutionary differences: we are most likely to focus on individual loci with large shifts in expression. However, even small changes in expression of one gene can be amplified via downstream effects of entire networks of genes, thereby exerting large phenotypic effects. Scanning large expression networks for correlated changes holds a great promise for uncovering evolving genes whose expression is either highly noisy with respect to genotype, or whose expression is only moderately shifted across populations.

Taken together, our snQTL analysis offers a powerful, effective, and adaptable framework for mapping QTLs that affecting network-based co-expression. We believe our approach brings a broad impact to the genetics community.

## 4 Materials and Methods

### Sparse matrix decomposition

We use penalized matrix decomposition [44] to approximately solve for sparse symmetric matrix decomposition in (4). The PDM with input matrix *D* is expressed as

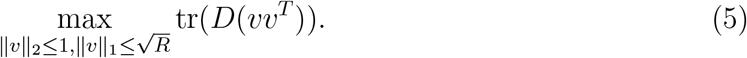

By [44], the solutions to (5) always have ∥*v*∥_2_ = 1 and satisfy the inequality ∥*v*∥^2^ ≤ ∥*v*∥_0_ ≤ *R*. Therefore, (5) is an good approximation to sLME in (4). We follow the algorithm in [44] to solve (5).

### Sparse symmetric tensor decomposition algorithm

We solve the optimization problem (2) via SSTD by an iterative algorithm. For a tensor 𝒟 ∈ ℝ^*p*1×*p*2×*p*3^ and vectors *v*^(*k*)^ ∈ ℝ^*pk*^ for *k* = 1, 2, 3, we define the tensor-by-vector product on mode 1, mode 2, and mode 3 as

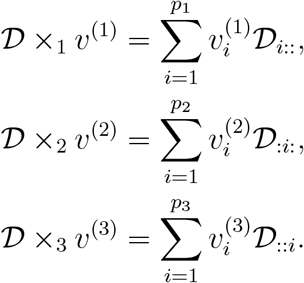

Given input tensor 𝒟, our decomposition algorithm is presented as follows:

1. *Input*. Differential tensor 𝒟 ∈ ℝ^*p*×*p*×3^, sparsity parameter *R*, and iteration number *T*.
2. *Initialization*. Randomly initilize the unit vectors *v*^(0)^ ∈ ℝ^*p*^, *u*^(0)^ ∈ ℝ^3^.
3. *For iteration t* = 1, …, *T*, alternatively update the decomposition components *v*^(*t*)^ and *u*^(*t*)^:

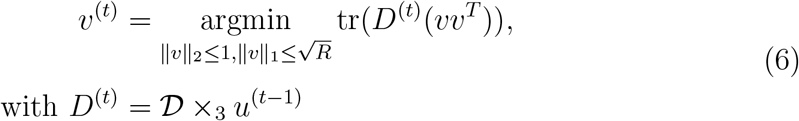

and

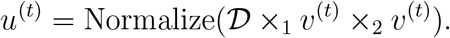
4. *Output*. Output the eigen components *v*(𝒟) = *v*^(*T*)^, *u*(𝒟) = *u*^(*T*)^ and estimated sLTE

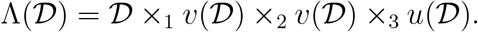

Here Normalize(*v*) = *v/*∥*v*∥_2_ denotes the vector normalization step. We make two comments on our algorithm. Previous work [28] enforces by value truncation. In contrast, our approach achieves sparsity through an optimization process called PMD (Proximal Minimization with Duality) during the update of a variable *v*^(*t*)^ in (6). Our approach is computationally faster and reflects the symmetry in our SSTD model. Second, in our construction of differential tensor input 𝒟, the third slide *D*_*BH*_ can be expressed as the sum of first two slides *D*_*AB*_ and *D*_*AH*_. While this linear relationship does not affect the final results of association testing, we choose to analyze the model using a full 3-layer tensor 𝒟 for easier interpretation.

### Permutation and empirical p-values

We used permutation to obtain empirical p-values based on our proposed test statistics. Specifically, at each marker, we repetitively shuffle three genotypes of samples, re-divide the expression dataset into three groups, and re-calculate the test statistics for *B* times. Let *S* denote the test statistic with original genotype, and *S*_*b*_ denote the test statistic with shuffled dataset in the *b*-th permutation for *b* = 1, …, *B*. We obtain the empirical p-value as

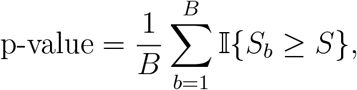

where 𝕀{·} is the indicator function.

In our stickleback data analysis, we first obtained the empirical p-values for all markers with *B*_0_ = 100 permutations for preliminary nQTL screening. For the markers showing preliminary empirical p-values smaller than 0.05, we re-ran the tests with *B* = 500 permutations for accurate p-values estimations.

### Sparsity hyperparameter

We set the sparsity parameter *R* to 0.25*p* for simulations and 0.09*p* for the stickleback data analysis. This aligns with the expectation that only a few thousand genes contribute to the main co-expression differences. Users can adjust *R* for a sparser (lower R) or denser (higher R) network based on their specific needs.

## 5 Acknowledgements

This study is supported by NSF FAIN-2133740 (to D.B, M.W, J.W., and J.H.), NSF CAREER DMS-2141865 (to M.W.), NIH 1R01AI123659-01A1 (to D.B.), HHMI Early Career Scientist funding (to D.B.), and 1R35GM142891-01 (to J.W.).

## Supporting Information Text

### A Extra analyses of simulated data

#### A.1 Simulated data generation

Our simulated dataset consists of both genotypes and expressions of *p* genes for *n* samples, represented as two *n*-by-*p* matrices *G* and *E*, respectively. To simulate genotypes, we started with homozygous parents and simulated the genotypes for an F1 cross followed by an F2 intercross generation with random chromosomal crossing overs. For each F1 gamete, we simulated one recombination event per chromosome, randomly placed along the chromosome with a uniform distribution. For extra analysis in Section 1A.3, we repeated the breeding procedures to simulate genotypes for F3, F4, and F5 intercross generations, to mimic the process of generating fine-mapping populations. The simulated genotype matrices have entries with values 0, 1, or 2, where 0 and 2 refer to homozygous genotypes, and 1 refers to the heterozygous genotype. We use *i* for the index of gene and use *j* for the index of sample. Given the simulated genotype matrix *G*, we simulate baseline expression level (null data) for each gene in each sample, assuming no network effects. Specifically, we simulated the null expression value of the *i*-th gene for the *j*-th sample independently based on Possion distribution with scalar parameters (*μ, σ*_*α*_, *σ*_*β*_):

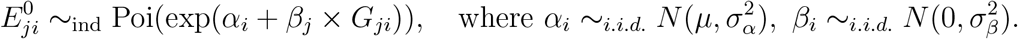

The parameters *μ* and 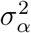 control the overall mean and deviation of the expression data, respectively, and 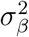 controls the degree of genetic effects on expression. Next, we constructed the gene coexpression networks, represented as symmetric matrices *M*_*k*_ ∈ ℝ^*p*×*p*^, *k* = 0, 1, 2, for three genotypes. For genotype *k*, the entries in *M*_*k*_ follow the following distribution independently with scalar parameters (*d*_*k*_, *δ*_*k*_):

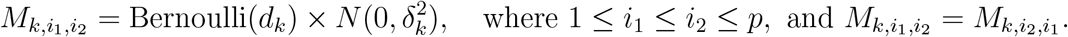

The parameter *d*_*k*_ controls the sparsity of gene-gene correlation (i.e. number of nonzero entries in *M*_*k*_), and 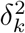 controls the magnitude of nonzero entries in *M*_*k*_ in genotype *k*. The genetic-related network effects are specified by setting various values of 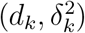. Furthermore, to mimic the additive genetic effects, we may set the heterozygous network as the average of homozygous networks, i.e., *M*_1_ = (*M*_0_ + *M*_2_)*/*2. For simplicity we do not consider more complex patterns of dominance or over-/under-dominance, which may exsit in empirical coexpression networks in hybrid populations.

Last, we imposed the coexpression network effect on top of the the null expression. We randomly selected one gene, say *i*^*\**^, as the nQTL. The genotype of nQTL *i*^*\**^ determined which matrix *M*_*k*_ is for the coexpression network effect. We generated the final expression data for genes involved in coexpressed pairs, using the following calculations:

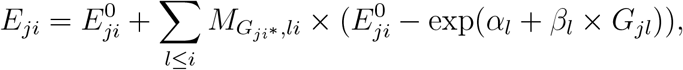

where the second term on the right-hand side is the network effect of nQTL *i*^*\**^ to the expression of gene *i* in sample *j*.

In our simulation, we generated data of *p* = 200 genes located on 20 chromosomes with varying population size *n* from 50 to 500. We considered additive network effect and tuned the parameters to mimic the distribution of expression levels from the empirical stickleback dataset, by choosing:

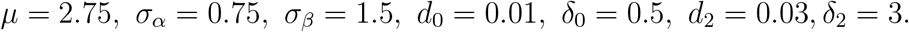

#### A.2 Raw correlation map for simulated data

Our analysis compared the absolute genetic correlation heatmaps between real stickleback data [35] and synthetic data (Figure S1). Unlike Figure 2A (real data), the synthetic heatmap shows correlations ranging from −1 to 1, while real marker pairs mostly have positive correlations. This difference arises from three aspects of our simulation design:

- Parent Genotype Initialization: In the simulation, half of first-generation diploid parents are given genotype AA while the other half are given genotype BB, for all markers. This leads to all offspring in the first generation having the same genotype AB for all markers.
- Enforced Crossover: During gamete formation, we simulate chromosomal crossover events. While these crossovers are important for real breeding, they can create negative correlations in our simulated data. This is because markers near the head and tail of the chromosome are likely to have different genotypes in the resulting offspring, even though they came from the same parent.
- Limited Breeding Generations: We only simulated breeding for two generations (F1 and F2). With more generations, these negative correlations due to initial crossovers would eventually be diluted through further recombination.

**Figure S1:**
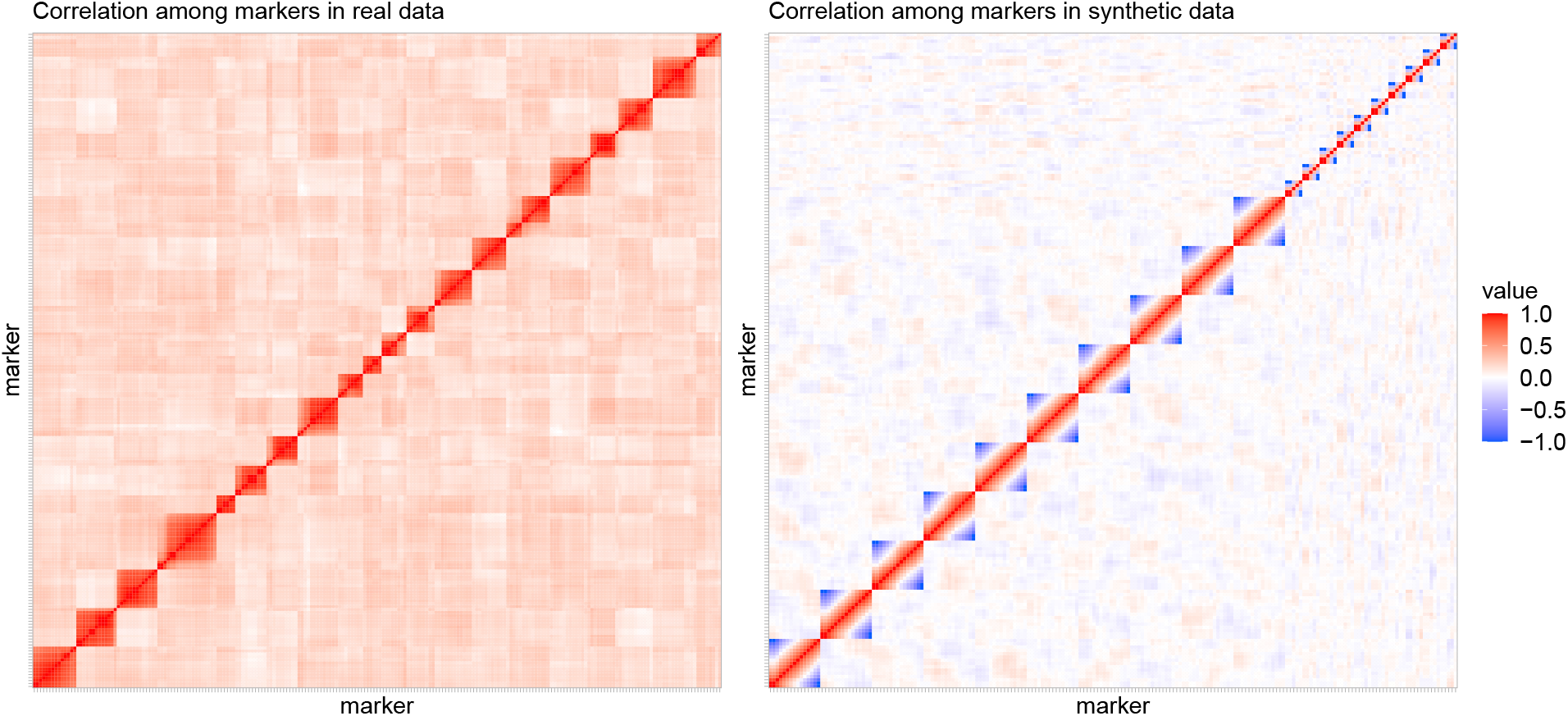
Genetic correlation heatmap among markers in real F2 hybrid three-spined stickleback data and synthetic data. Blue color indicates negative correlation; red color indicates positive correlation.

For example, consider a chromosome with 5 markers, with 0 representing genotype A and 1 representing genotype B. The second-generation diploid parents have two chromatids for chromosome 1, with genotype (00000) and (11111). With a crossover at the fourth marker, possible gametes could be (00011) and (11100). Combine two random gametes to obtain the genotype of diploid offspring. If marker 1 has genotype (0,0), then marker 5 must have (1,1); if marker 1 has genotype (0,1), then marker 5 must have (1,0). Markers at opposite ends of the chromosome have opposite genotypes, leading to a negative correlation. It is important to note that our snQTL testing is invariant to the label switch between A and B. Therefore, the presence of negative correlations in the simulated data does not impact the evaluation of our method’s effectiveness.

#### A.3 Simulation for synthetic data with more generations

Figure S2 presents additional simulation results using synthetic data generated from F3, F4, and F5 hybrid crosses. These later breeding generations are often used in fine-mapping studies because they create smaller linkage windows. In essence, these later generations move closer to mimicking GWAS in genetically diverse outbred populations, where breeding is not involved. We find that our comparison conclusions remain the same to those performed with the F2 hybrid data in the main text. This consistency demonstrates that snQTL maintains its outer-performance compared to the local method, regardless of the breeding generation used in the simulation.

### B Pre-processing and extra analyses with stickleback data

#### B.1 Pre-processing of stickleback data

The original stickleback data Weber et al. [35] includes gene expression levels (transcript counts) for 26,285 genes and genotypes for 234 genetic markers from 351 samples in the F2 and backcross generations. The data also contains covariate information such as sex, ancestry, and infection status.

We describe our pre-processing on the gene expression matrix. Let *X*^0^ denote the raw transcript count matrix, and *n* = 351, *p* = 26, 285, *q* = 234 for the numbers of samples, genes, and markers, respectively We use *i* for the index of gene and use *j* for the index of fish.

We follow these three main steps:

i. Normalization. We normalized the raw transcript counts to account for differences in sequencing depth across samples. This is achieved by dividing each transcript count 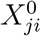 by the total transcript count for that sample. The resulting normalized matrix is denoted by *X*^*N*^ with entries

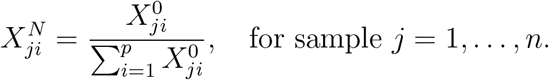

The denominator 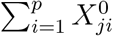 is referred to as the library size or sequencing depth of the sample *j*.
ii. Covariate Removal. We removed the effects of covariates of sex and ancestry. This was done using linear regression. The covariate sex has three levels (female, male, and non-identified), and the covariate ancestor has 36 levels based on the parental origin. The residuals from this model represent the gene expression levels after removing the influence of these covariates.

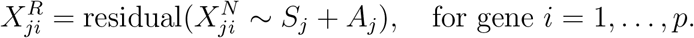

Here, *y ∼ x* represents the standard repression model in which *y* servers as the response and *x* serves as predictor, and *S*_*j*_ denotes the sex effect and *A*_*j*_ denotes the ancestor effect. We use *X*^*R*^ to denote the residual matrix after removing covariate effects. As we mentioned in the main text, we also consider warm infection status as a possible covariate. We explored this further in Section 2 B.3 using a similar regression approach with an additional binary predictor for worm presence/absence:

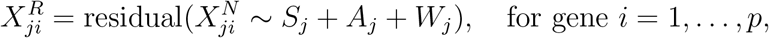

where *W*_*j*_ is a binary predictor encoding the worm presence/absence of the samples.
iii. Gene Selection: Finally, we selected the top 10,000 genes with the highest adjusted mean expression *m*_*i*_ defined by

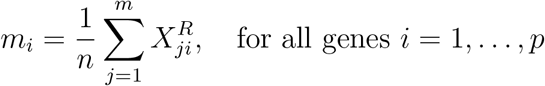

This metric represents the average expression level for each gene across all samples. We focused on highly expressed genes because they are more likely to be relevant for biological processes. The cutoff of 10,000 genes was chosen for computational efficiency.

**Figure S2:**
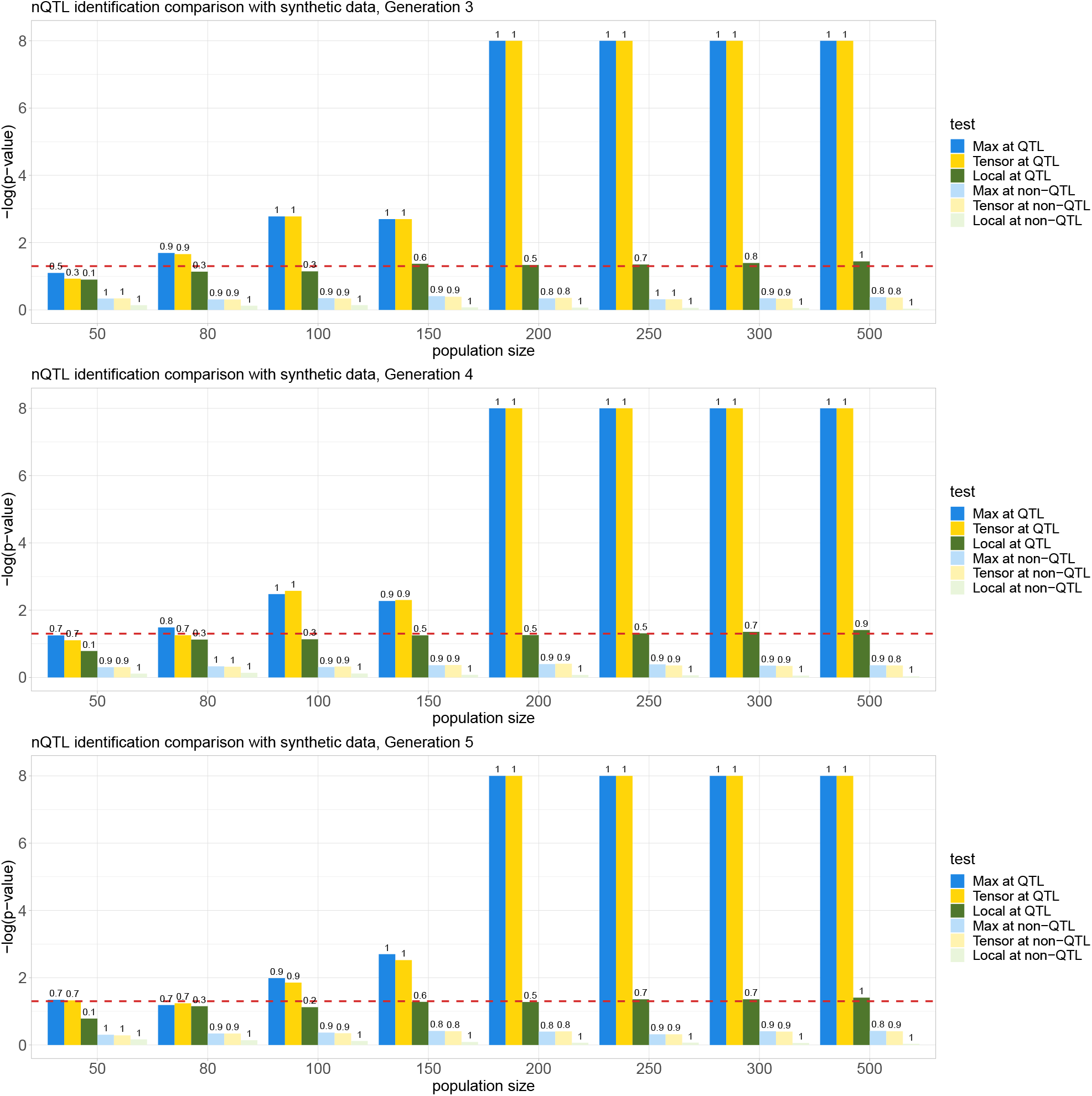
nQTL comparision for snQTL framework and local method (F-test for regression of pairwise co-expression against genotype) on synthetic data with varying population size from 50 to 500. Synthetic datasets are generated from the F3, F4, and F5 hybrids (from top to bottom), respectively. Red dashed line corresponds to the critical threshold with p-value 0.05. True positive (or negative) rates for the tests at nQTL (or non-nQTL) are shown above the bars. All reported numbers are averaged across 15 replications for each population size.

#### B.2 Testing results with matrix statistics

In addition to the tensor statistics, we investigated two alternative approaches based on matrix-spectrum statistics. The first one uses the max statistic defined in the main text. The second approach utilizes a variant called the sum statistic, defined as:

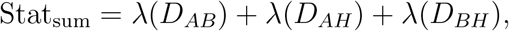

where *λ*(·) represents the sLME, *D*_*AB*_, *D*_*AH*_, and *D*_*BH*_ represent pairwise differential network from the original sample correlation matrices.

As shown in Figures S3 and S4, we found that both the max and sum statistics identified nQTLs for sticklebacks clustered on chromosomes 3, 8, and 18. This consistency across different statistical methods strengthens the reliability of our findings.

**Figure S3:**
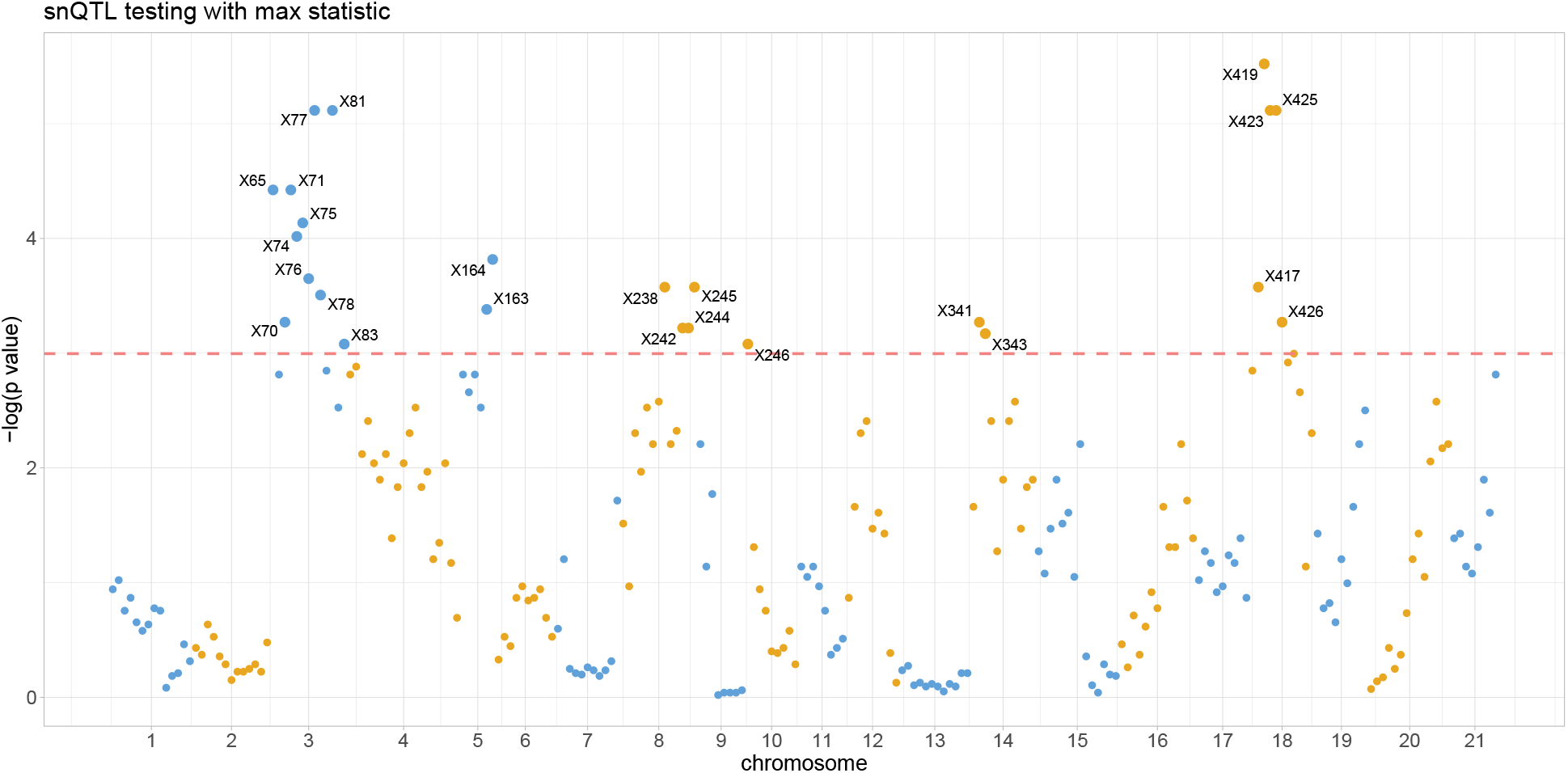
Manhattan plot for snQTL testing with max statistics marks stickleback nQTLs (above pink dashed line with p-values smaller than 0.05), mainly clustered in Chr3, Chr8, and Chr18.

#### B.3 Testing results controlling for tape worm infection

To prioritize analyses with limited computational resources, we performed snQTL testing only on the top nQTLs identified from the non-infection-controlled expression data. We found that the top nQTLs on chromosomes 3, 8, and 18 remained the same even after controlling for infection status (Table S1). This consistency suggests that the network effects of these nQTLs are not driven by the environmental factor of worm infection. Rerunning the analysis with infection-controlled expression data for all markers might reveal even more significant nQTLs and related discoveries. However, we will leave such additional analyses for future studies.

**Figure S4:**
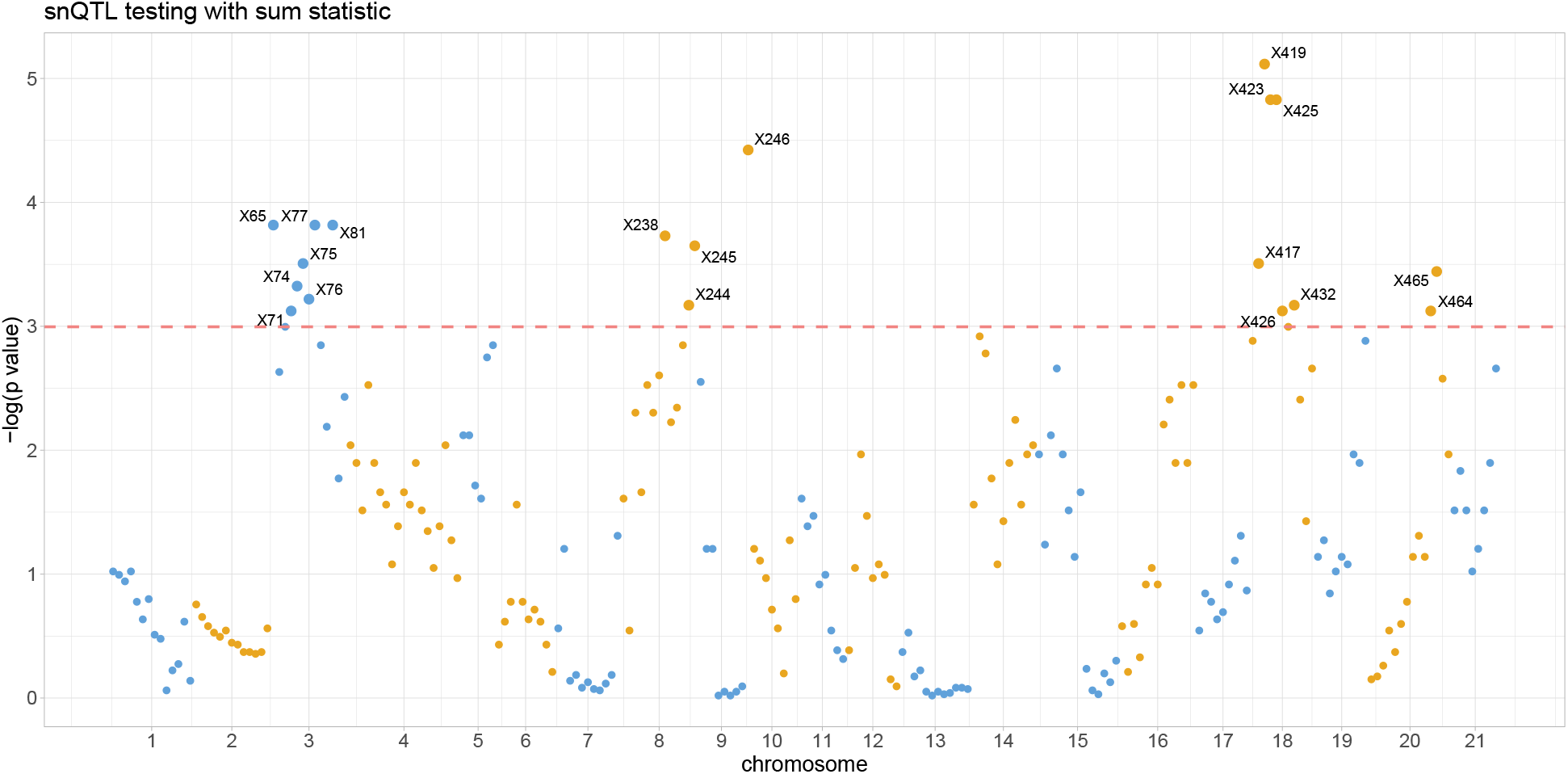
Manhattan plot for snQTL testing with sum statistics marks stickleback nQTLs (above pink dashed line with p-values smaller than 0.05), mainly clustered in Chr3, Chr8, and Chr18.

**Table S1:** Empirical p-values of top nQTLs obtained by snQTL testing with infection-uncontrolled expression (main text result) and with the infection-controlled expression (Supporting Information Text Section B.3).

| Markers | p-values (infection uncontrolled) | p-values (infection controlled) |
| --- | --- | --- |
| X77 | 0.01 | 0.002 |
| X81 | 0.012 | 0.004 |
| X419 | 0.008 | 0.004 |
| X423 | 0.008 | 0.008 |
| X238 | 0.018 | 0.024 |
| X245 | 0.02 | 0.016 |

#### B.4 Population branch statistic distributions on Chr 3 and Chr 8

Figures S5 and S6 show that several protein-coding genes lie in regions adjacent to PBS outliers near the nQTLs on Chr 3 and Chr 8.

**Figure S5:**
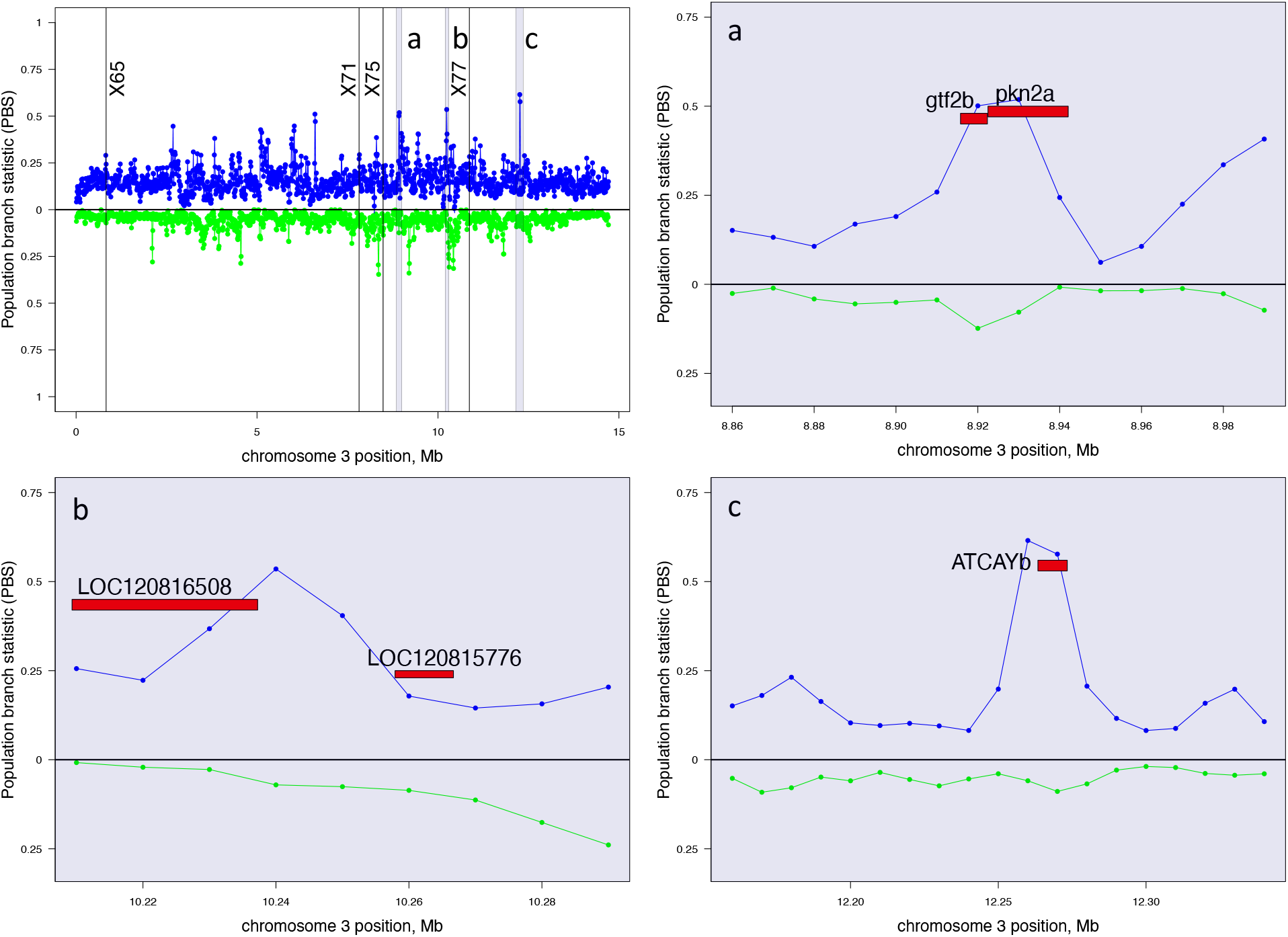
Strong genomic targets of selection with high population branch statistic distribute around the outstanding nQTLs in Chr 3. Values above the medial line represent higher PBS in Gosling Lake (blue); values below the line represent higher PBS in Roberts Lake (green). Protein-coding genes lie in regions adjacent to three PBS outliers (a, b, c) around nQTLs (markers X65, X71, X75, and X77).

**Figure S6:**
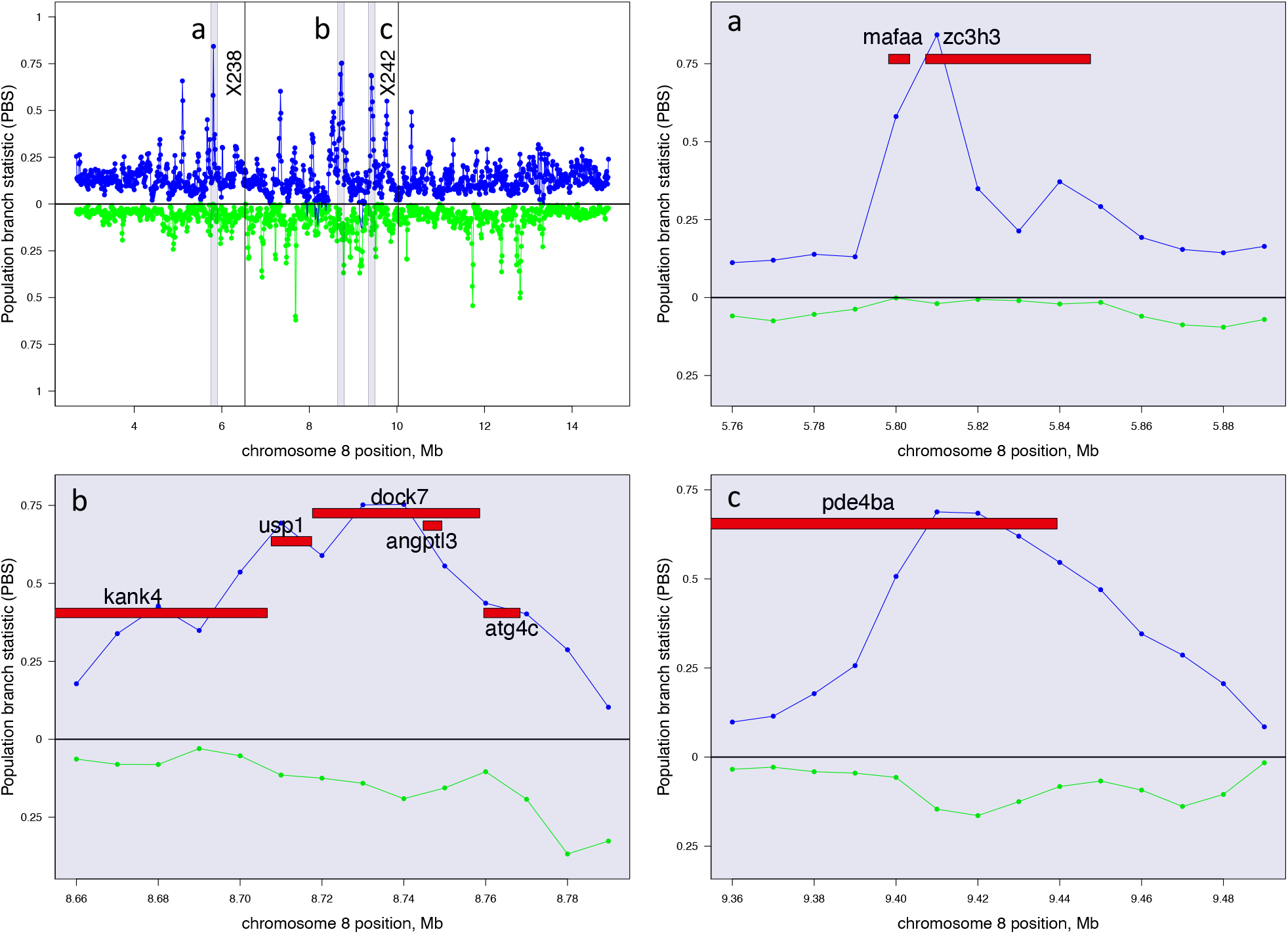
Strong genomic targets of selection with high population branch statistic distribute around the outstanding nQTLs in Chr 8. Values above the medial line represent higher PBS in Gosling Lake (blue); values below the line represent higher PBS in Roberts Lake (green). Protein-coding genes lie in regions adjacent to three PBS outliers (a, b, c) around nQTLs (markers X238 and X242).

#### B.5 Joint differential networks for nQTLs on Chr 3 and Chr 8

We further analyzed the top nQTLs identified earlier: X77 on Chr 3 and X238 on Chr 8. Interestingly, the results at these loci were highly similar to those observed at X419 on Chr 18 (see main text). Figure S7 illustrates this similarity. The sets of top 100 genes identified by high tensor leverages at X77 and X238 show a significant overlap with the top genes for X419, especially the top 10 most important genes. However, the specific ranking of these genes might differ slightly between nQTLs. To explore this further, we constructed a joint differential network for X77 and X238, using the top genes identified at X419. Figure S8 reveals the strong connections between primary and secondary genes. This patten is consistent across these multiple nQTLs. This consistency strengthens the evidence that our findings regarding oxygen transport pathways in the joint differential network are not simply random observations.

**Figure S7:**
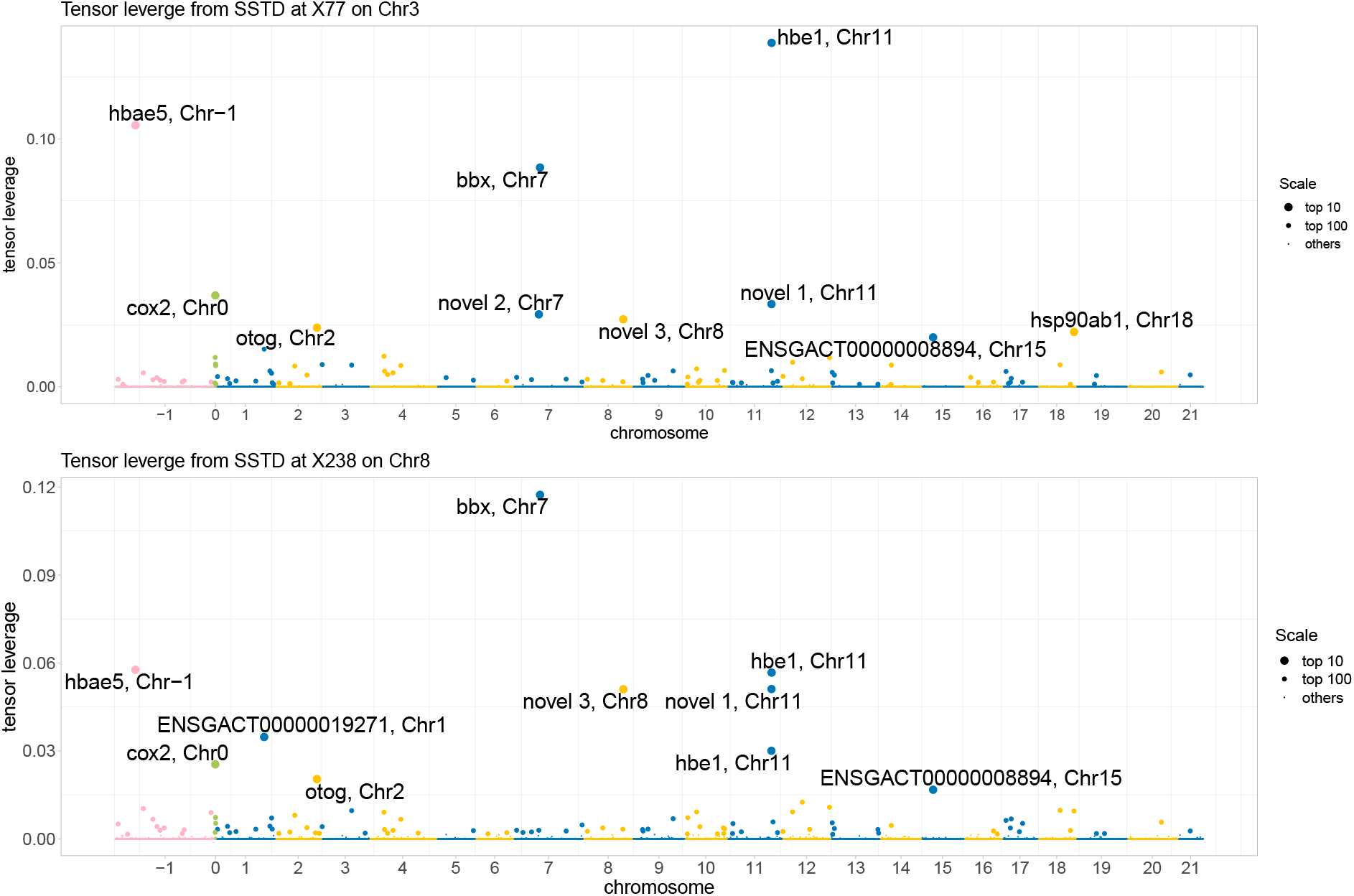
Joint differential network analysis at nQTLs X77 on Chr 3 (top) and X238 on Chr 8 (bottom) with leverage scores for 10000 genes. Primary genes with top 10 leverage are highlighted with transcription IDs. Mitochondrial genome (MT) and scaffold region are coded as Chr 0 and Chr −1, respectively.

**Figure S8:**
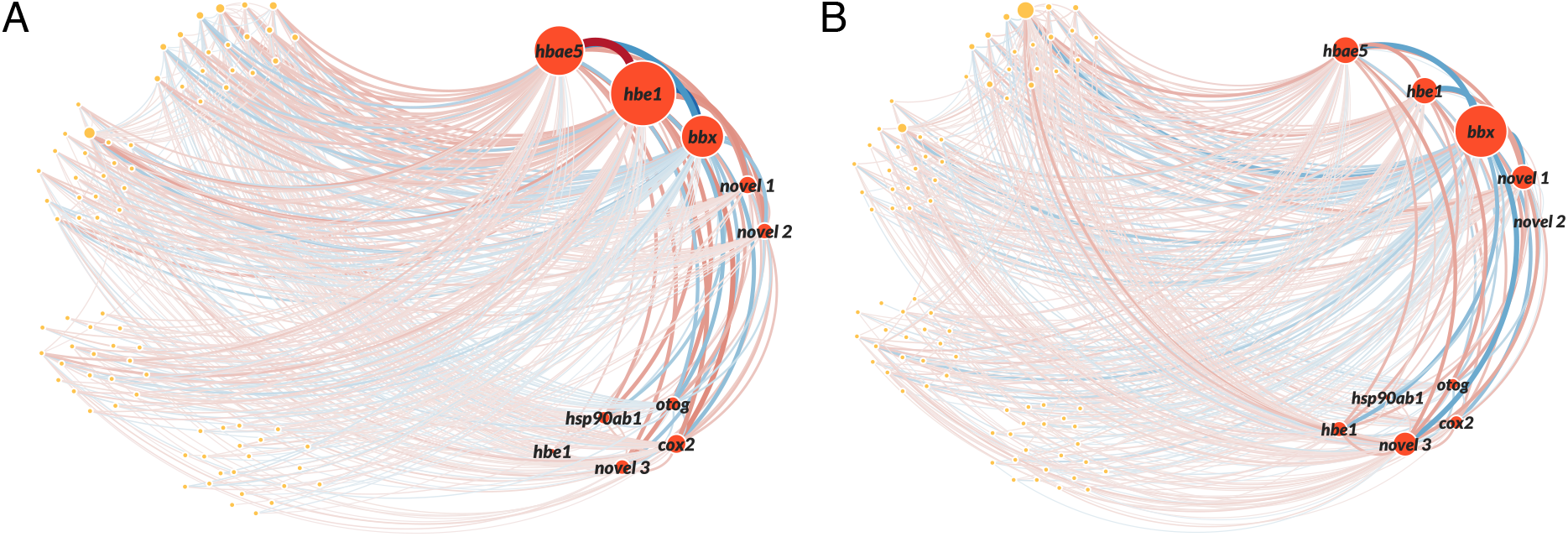
Joint differential networks at nQTLs (A) X77 on Chr 3 and (B) X238 on Chr 8 with top 100 genes identified in the analysis at X419 on Chr 18. The edge width indicates the connection strength between two genes; the diameter of node indicates the leverage of the genes; the color indicates the enhancement (red) or reduction (blue) of the connection compared with average level. Top 10% strongly connected edges are selected. novel 1: ENSGACT00000018413; novel 2: ENSGACT00000026589; novel 3: ENSGACT00000017116.

